# Plasma regulates homeostatic pulmonary endothelial signaling to mitigate vascular leak following polytrauma and hemorrhagic shock

**DOI:** 10.64898/2026.03.04.709561

**Authors:** Taylor E. Wallen, Krystal L. Rivera-Figueroa, James D. Odum, Giacynta Vollmer, Lei Zheng, Melad N. Dababneh, Delores A. Stacks, Camilla Margaroli, Robert P. Richter, Jillian R. Richter

## Abstract

**Introduction:** In hemorrhagic shock, plasma resuscitation preserves vascular integrity and protects against trauma-induced coagulopathy and organ injury. Despite demonstrated clinical benefit, the endothelial mechanisms underlying plasma resuscitation remain incompletely defined. This study investigated endothelial-specific responses to plasma resuscitation to identify targetable pathways that promote vascular repair after traumatic injury.

**Methods:** A murine model of severe polytrauma-hemorrhagic shock (PT/HS) with demonstrable vascular endotheliopathy by 24 hours was used to compare pulmonary vascular endothelial cell (EC) responses to resuscitation with lactated Ringer’s (LR) relative to fresh frozen plasma (FFP). Whole blood was collected for inflammatory biomarker analysis, and pulmonary vascular leak was quantified by dextran extravasation. Pulmonary EC glycocalyx (eGC) structure was assessed by transmission electron microscopy and immunofluorescence. Spatial transcriptomic profiling of pulmonary ECs was performed using a GeoMx Digital Spatial Profiler. Key transcriptomic findings related to mitochondrial biogenesis were validated by immunostaining and by treating primary human lung EC with FFP or LR.

**Results:** At 24 hours after injury, FFP reduced systemic inflammatory cytokines, pulmonary innate immune cell infiltration, and PT/HS-induced vascular leak compared to LR. Plasma levels of syndecan-1, syndecan-4, and hyaluronan were decreased, consistent with enhanced pulmonary eGC expression. Although few differences in eGC-related genes were detected, pathway analysis revealed enrichment of cellular bioenergetics and metabolic recovery pathways in ECs after FFP, whereas LR was associated with oxidative stress and inflammatory signaling. FFP enhanced mitochondrial content in pulmonary EC after PT/HS and in treated human EC compared to LR-treated controls.

**Conclusions:** FFP resuscitation after PT/HS reduces systemic inflammation and preserves pulmonary vascular barrier function, potentially through promotion of mitochondrial signaling, metabolic recovery, and endothelial stress regulation.

## Introduction

Trauma-related injuries are the leading cause of death in individuals under 45 and remain a global health burden, contributing to over 5.8 million deaths annually.^1,2^ While advances in trauma care have reduced early mortality from hemorrhagic shock, clinical complications—mainly related to multiple organ injury (MOI)—persist during hospitalization, accounting for up to 25% of trauma-related fatalities.^3–7^ A central, unifying driver of trauma-induced MOI is vascular dysfunction, rooted in the rapid degradation of the endothelial glycocalyx (eGC).

The eGC is a protective, matrix-like barrier on the luminal surface of endothelial cells (EC), composed primarily of heparan sulfate proteoglycans (HSPG) and glycosaminoglycans (GAGs) that are essential for maintaining EC homeostasis and monolayer integrity, regulating vascular permeability, and balancing thrombo-inflammatory responses.^8^ Within minutes of traumatic injury, inflammatory mediators and enzymes damage the eGC, initiating a cascade of signaling events that disrupt EC homeostasis. Vascular endotheliopathy, in turn, promotes vascular leakage, EC inflammation, and coagulopathy that impairs microvascular (MV) perfusion and culminates in MOI.^9–11^

Plasma transfusion has been shown to protect the eGC from degradation, promote eGC restoration, and attenuate EC dysfunction in preclinical models of trauma and hemorrhagic shock.^12,13^ Clinically, early plasma administration in hemorrhaging patients helps mitigate coagulopathy, reduce transfusion requirements, and improve mortality.^14–16^ However, despite the recognized benefits of plasma administration in traumatically injured patients, patient response to plasma is heterogenous,^17^ and our understanding of the mechanisms by which plasma protects the injured endothelium remains limited. Moreover, due to persistent strains on available donor blood products,^18^ there is a critical need to better understand how plasma regulates eGC integrity and vascular functions that could inform the development of new adjunctive therapies to mitigate systemic endotheliopathy and organ injury after trauma. The goal of this study was to elucidate endothelial-specific mechanistic responses to plasma resuscitation, which may enable the identification of targetable pathways for promoting vascular repair and improving patient recovery following traumatic injury.

## Materials and Methods

### Animal Model of Trauma and Hemorrhagic Shock

Animal procedures were conducted under a protocol approved by the University of Alabama at Birmingham Institutional Animal Care and Use Committee and in accordance with Animal Research: Reporting of *In Vivo* Experiments (ARRIVE) guidelines. Twelve to fourteen-week-old adult male C57Bl/6 mice (Charles River) weighing 25-30 grams were allowed at least 72 hours to acclimatize to the environment with access to food and water ad libitum under 12-hour light/dark cycles prior to experimental use.

A murine model of polytrauma-hemorrhagic shock (PT/HS) was utilized as previously described.^19–22^ Mice were anesthetized by intraperitoneal injection of pentobarbital (70 mg/kg). The abdomen and groins were shaved and washed with 10% povidone-iodine. A 2-centimeter midline laparotomy was performed to induce soft tissue trauma followed by cannulation of both femoral arteries to enable continuous monitoring of systemic arterial pressure and simultaneous vascular access for hemorrhage and resuscitation. After a 10-minute stabilization period, blood was removed via the femoral arterial catheter to achieve a mean arterial pressure (MAP) of 25 ± 5 mmHg, which was maintained for 60 minutes by additional bleeding if required. After the 60-minute hemorrhagic shock period, mice were resuscitated over 20 minutes with either lactated Ringer’s solution (LR) at a volume equivalent to three-times the shed blood volume or with fresh frozen plasma (FFP) to achieve a target MAP of 70-80 mmHg, representing fixed-volume crystalloid versus modern goal-directed blood product resuscitation, respectively. FFP was prepared from age-matched, male C57Bl/6 mice by collecting whole blood via cardiac puncture, anticoagulating the blood using citrate-phosphate-double dextrose (CP2D), and centrifuging anticoagulated blood at 1500 *x g* for 15 minutes. Plasma was collected, centrifuged again to remove residual blood cells, and stored in single-use aliquots at −20°C until use. Following resuscitation, bilateral pseudo-fractures and muscle crush injuries were performed to induce polytraumatic injuries via injection of 200 µL of sterile bone homogenate, collected from littermates of the injured mouse, into both hindlimbs followed by a 30-second, bilateral clamping of the hamstring muscles. Sham mice underwent femoral artery cannulation for pressure monitoring but were not subjected to hemorrhage or other tissue injuries. Sustained-release buprenorphine (0.05 mg/kg) was administered subcutaneously as analgesia following resuscitation. Mice were monitored for 24 hours after resuscitation and anesthetized with intraperitoneal 80 mg/kg ketamine and 10 mg/kg xylazine prior to blood and tissue harvest.

### Complete Blood Cell Counts and Plasma Biomarker Analyses

Murine blood was collected via cardiac puncture, and complete blood counts from whole blood were obtained using a HemaVet 950FS (Drew Scientific). Plasma was generated from whole blood by centrifugation at 1,500 *x g* for 15 minutes at 20-22°C and stored at −80°C for future analyses. Plasma levels of inflammatory cytokine and chemokine were assessed using a customized Luminex Mouse Discovery 15-Plex Assay (R&D Systems) to measure the following: angiopoietin-2 (Angpt-2), CXCL2, soluble intercellular adhesion molecule (ICAM)-1, interferon gamma (IFNγ, interleukin (IL)-1a, IL-1b, IL-6, IL-10, keratinocyte-derived chemokine [KC; murine ortholog of human chemokine (C-X-C motif) ligand 1, or CXCL1], macrophage inflammatory protein (MIP)-1a, MIP-2a (murine ortholog of human CXCL2), regulated upon activation, normal T expressed and secreted (RANTES), syndecan-1 (Sdc1), tumor necrosis factor alpha (TNFα), vascular endothelial growth factor (VEGF)-A, and soluble VEGF receptor (VEGFR)-2. Commercially available enzyme-linked immunosorbent assays (ELISA) for mouse glypican-1 (Gpc1) (Biomatik), hyaluronan (HA) (Echelon Biosciences), and syndecan-4 (Sdc4) (Novus Biologicals) were performed according to manufacturer specifications.

### Lung Histology and Immunostaining

Lung tissues were fixed in 10% neutral buffered formalin (NBF) for 24 hours, followed by transfer to 70% histological-grade ethanol. Fixed lung tissues were then paraffin-embedded and sectioned. Sections for histological evaluation and immunohistochemistry were stained with hematoxylin and eosin (H&E). The presence of acute lung injury was assessed using H&E sections imaged under a 40X objective by a board-certified pathologist who was blinded to the treatment group. Lung injury was scored using a 4-point scale, as previously described. ^23–25^

Immunohistochemistry was performed using H&E-stained sections following deparaffinization, rehydration, and antigen retrieval. Tissues were blocked with 1% bovine serum albumin and 10% normal animal serum, and polymorphonuclear (PMN) inflammatory cells were identified using a monoclonal rat anti-mouse Ly-6B.2 alloantigen antibody (Bio-Rad Laboratories; 1:500 v/v in blocking buffer), and macrophages were identified using a monoclonal rat anti-mouse F4/80 antibody (eBioscience; 1:50 v/v in blocking buffer). Mitochondrial content was assessed using recombinant rabbit translocase of outer membrane mitochondria 20 (TOM20) antibody (Thermo Fisher Scientific; 1:200 v/v in blocking buffer).

Primary antibodies were detected using a rabbit anti-rat biotinylated antibody (Vector; 1:200 in 0.1% TBS-T) followed by incubation with horseradish peroxidase (HRP)-labeled streptavidin (BD Pharmingen). HRP activity was detected using an UltraVision LP Detection System: HRP Polymer & DAB Plus Chromogen kit. Total cells and positively stained cells were identified and quantified using the CellQuant method within QuantCenter (3DHISTECH, version 3.0). The software applies pre-specified algorithms for tissue pre-segmentation, nuclei detection (size 5, counterstain), and automated exclusion of folding artifacts, to objectively determine the total cell (nuclei) counts, number of positively-stained cells and the histoscore (H-score)—a semi-quantitative metric that combines the staining intensity with the percentage of positive cells. PMN and macrophages counts were determined in entire lung tissue sections, and TOM20 staining was quantified by averaging the H-score of five vascular regions of interest per tissue.

### Vascular Permeability Assay

Vascular permeability was assessed 24 hours after PT/HS by quantifying the extravasation of fluorescently labeled dextran into pulmonary tissue, as previously described.^26,27^ Briefly, 200 μL of 0.2 mg/mL Alexa Flour (AF) 680-labeled dextran (10 kDa; Thermo Fisher Scientific) was administered via retroorbital injection into mice anesthetized using intraperitoneal 80 mg/kg ketamine and 10 mg/kg xylazine. After 30 minutes, mice were perfused with cold PBS via the left ventricle to remove intravascular tracer, and lungs were subsequently harvested and imaged using the LI-COR Odyssey imaging system (LI-COR Biosciences, Lincoln, NE). Image Studio software (LI-COR Biosciences) was used for image analysis and quantification of average fluorescence signal intensity per area within each lung.

### eGC Immunofluorescence Staining

The pulmonary vasculature was flushed to remove residual blood cells by cannulating the pulmonary artery and perfusing cold saline for 3 minutes at a pressure of 12 cm H_2_O. After flushing, lungs were perfused with 10% NBF using a syringe pump at a rate of 0.25 mL/min for 30 minutes to initiate tissue fixation. Lungs were then excised, fixed in 10% NBF for 24 hours and dehydrated as above. Fixed lung tissue was then paraffin-embedded and sectioned for histological analysis. For eGC staining, tissue sections were blocked with 1% bovine serum albumin (BSA) for 1 hour, then incubated with a polyclonal goat anti-mouse CD31/PECAM-1 antibody (1:200; R&D Systems) to label ECs. Following incubation with a rabbit anti-goat AF-488 secondary antibody (1:500; Invitrogen), sections were stained with AF594–conjugated wheat germ agglutinin (WGA; 1:1000; Thermo Fisher Scientific) for 1 hour to detect N-acetylglucosamine and sialic acid residues within the eGC. To minimize background signal, the trueVIEW™ Autofluorescence Quenching Kit (Vector Laboratories) was applied, followed by nuclear staining with DAPI (1:1,000; Thermo Fisher Scientific). Fluorescent images were acquired using a Crest X-Light V3 spinning disk confocal microscope (Nikon) and analyzed with NIS Elements software (Nikon).

### Transmission Electron Microscopy

To visualize the endothelial-specific glycocalyx layer in the lung microvasculature using electron microscopy, anesthetized mice were perfused through a cannula inserted into the pulmonary artery with a fixation solution containing 2% glutaraldehyde, 2% sucrose, 0.1 M cacodylate buffer (pH 7.3), and 2% lanthanum nitrate. To confine perfusion to the pulmonary circulation, small incisions were made in the inferior vena cava and left ventricle. The fixation solution was delivered using a syringe pump at a rate of 0.5 mL/min for 30 minutes. Following perfusion, the lungs were excised, cut into ∼1 mm³ pieces, and immersed in the same fixation solution for 2 hours at 4 °C. After primary fixation, tissues were transferred to a staining solution composed of 2% sucrose, 0.1 M cacodylate buffer (pH 7.3), and 2% lanthanum nitrate, and incubated overnight at 4 °C. Tissues were then washed twice in an alkaline sucrose solution (0.03 M NaOH in 2% sucrose), osmicated with 0.25% osmium tetroxide, and dehydrated through a graded ethanol series. Samples were embedded in epoxy resin, sectioned into ultrathin slices (90 nm), and imaged using a JEM-1400 High Contrast TEM (JEOL Ltd.) equipped with a NanoSprint43L-MarkII camera (Advanced Microscopy Techniques).

### GeoMx Digital Spatial Profiling

Five-micrometer–thick unstained lung sections designated for GeoMx digital spatial profiling (four sections per sham, LR, and FFP groups) were deparaffinized, rehydrated, and subjected to antigen retrieval. Sections underwent 16-hour in situ hybridization at 37°C with the GeoMx Mouse Whole Transcriptomic Atlas RNA probe mix (Bruker Corporation), followed by stringent washing and blocking in Buffer W. Tissues were then incubated with fluorescently labeled antibodies to identify ECs, epithelial cells, smooth muscle, and nuclei, respectively: AF647–conjugated anti-CD31/PECAM-1 (Bio-Techne; 1:50), AF594–conjugated anti-EpCAM (Abcam; 1:82), AF532–conjugated anti–myosin heavy chain 11 (Abcam; 1:20), and SYTO13 (Invitrogen; 1:167). After imaging on the GeoMx platform (Bruker Corporation), vascular segments were defined by anatomical criteria:^28^ pulmonary arteries as medium-to-large vessels adjacent to airways within bronchovascular bundles; pulmonary veins as medium-to-large vessels separate from bronchovascular bundles or at the lung periphery; and pulmonary microvasculature as alveolar-only regions. Regions of interest (ROI) were selected from ≥3 representative sections per vascular segment. RNA probes were released from ECs within each ROI by UV exposure and sequenced. Pre-specified GeoMx DSP quality control metrics were applied to raw transcriptomic data, followed by upper-quartile (Q3) normalization of ROI counts. Final normalized counts were uploaded to Mendeley Data and are publicly available here: doi: 10.17632/pskngd7fn8.1.

### Bioinformatic Analysis

Transcriptomic analysis was performed using RStudio in the R environment (version 2025.5.1+513). An unbiased approach was used at the outset to report the top 40 differentially expressed genes based on statistical thresholds of -log10 *p*-value > 1.3 and an absolute log2 fold-change > 0.5. *pheatmap* was used to generate the resulting heatmaps, *EnhancedVolcano* was used to generate volcano plots, and *ggplot2* was used to generate the bubble plot. Gene set enrichment analysis (GSEA) was performed using the ToppGene suite with Gene Ontology (GO): Biological Process and Reactome pathway datasets. The top GO: Biological Process and Reactome pathways were selected based on ≥5 gene overlap counts and false discovery rate (FDR)-adjusted *p*-value <0.05. We also took a targeted approach to evaluate genes associated with maintenance of eGC expression.

### In Vitro Treatment of Endothelial Cells

Primary human lung vascular ECs (Cell Applications) were seeded into Ibidi u-Slide VI 0.4 ibiTreat channels (Ibidi) pre-coated with attachment factor solution (Cell Applications) at 75,000 cells per channel. Confluent monolayers were treated with sterile LR or FFP (30% v/v in growth media) obtained from a healthy donor, as described previously.^29^ After 6 hours of treatment, cells were fixed with 4% paraformaldehyde for 15 min at 20-22°C, followed by immunostaining for TOM20 (1:200 v/v in PBS) to assess mitochondrial content and vascular endothelial (VE)-cadherin (eBioscience; 1:100 v/v in PBS) to assess cell-cell junctional integrity. Nuclear counterstain DAPI (Thermo Fisher Scientific; 1:1,000 v/v in PBS) was also applied. Fluorescence intensity of TOM20 expression was captured using a Nikon Crest spinning disk confocal microscope and quantified using ImageJ.

### Statistical Analysis

Data were assessed for normality using the Kolmogorov-Smirnov test. All data were normally distributed and, thus, presented as means with standard deviation. Statistical significance between sham and PT/HS treatment groups was determined using a one-way analysis of variance (ANOVA). For significant results (p<0.05), Fisher LSD post hoc test was employed given the low number of *a priori* pairwise comparisons and relatively small sample size within each group. A 2-sided α level of 0.05 was used to determine significance. Statistical comparisons were performed using GraphPad Prism, version 10.

## Results

### Plasma mitigates systemic inflammation and vascular leak after PT/HS

To recapitulate severe traumatic injury resulting in endothelial activation and vascular dysfunction, we employed a combined murine model of polytraumatic injuries and hemorrhagic shock, followed by fixed-volume resuscitation with LR or hemodynamically-guided resuscitation with FFP (**Figure 1A-C**). Twenty-four hours after resuscitation, PT/HS mice in both groups exhibited reduced white blood cell counts and significantly lower red cell blood counts and hematocrit levels compared to sham mice, though platelet counts were not significantly different (**Figure 1D-G**). LR-resuscitated mice had elevated levels of circulating inflammatory cytokines compared to sham controls and FFP mice, whereas plasma cytokine levels in mice resuscitated with FFP were mostly comparable to sham mice at 24 hours post-injury (**Figure 1H**). Furthermore, there were increased numbers of Ly6B^+^ PMN (**Figure 1I**) and F4/80^+^ macrophages (**Figure 1J**) in lungs of LR-resuscitated mice compared to sham and FFP groups, although no significant differences in histologic lung injury scores were observed between groups (**Supplemental Figure 1**).

**Figure 1.**
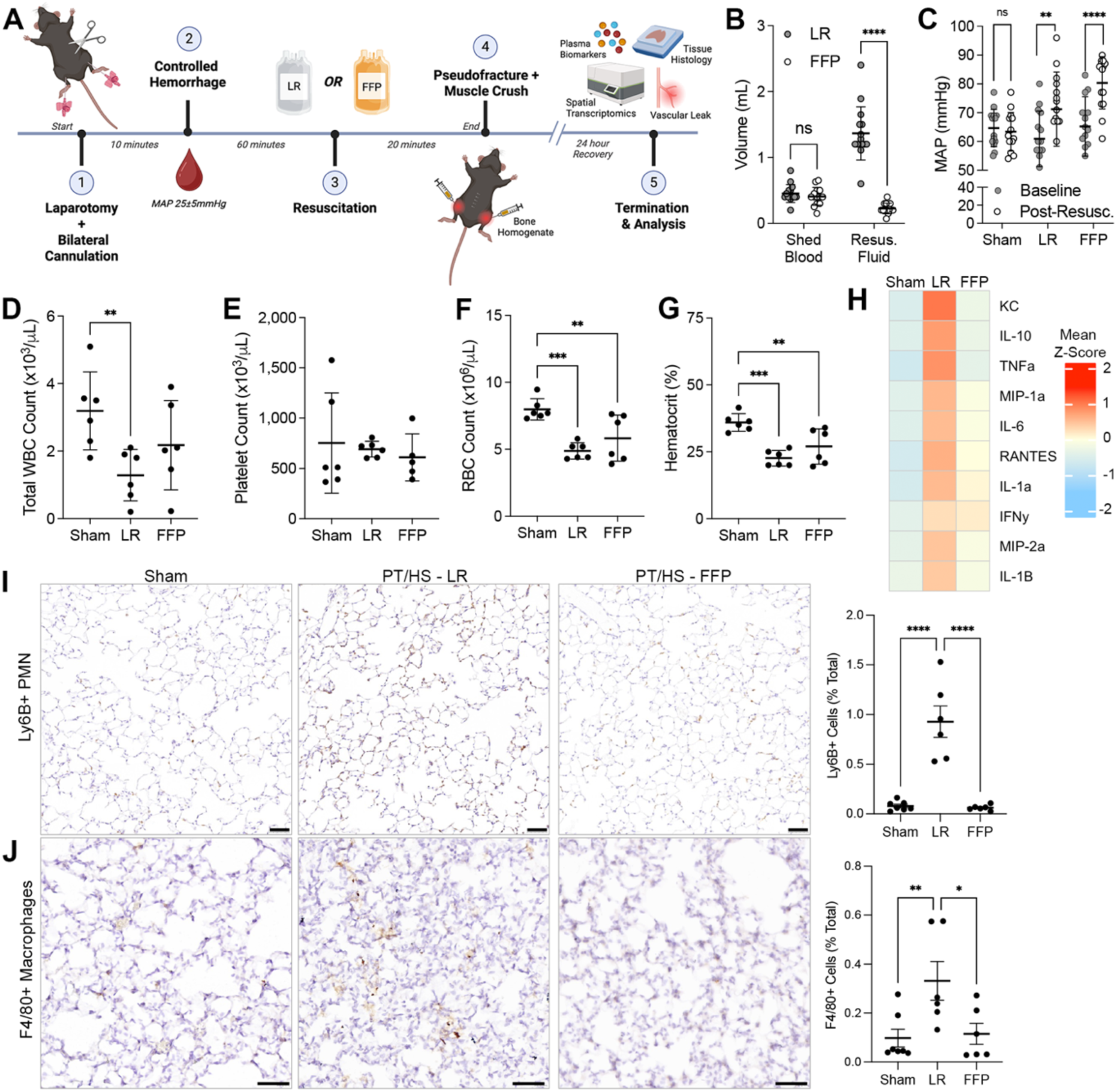
Administration of fresh frozen plasma (FFP) mitigates systemic and pulmonary inflammation after polytrauma and hemorrhagic shock (PT/HS). **(A)** Murine model of PT/HS with midline laparotomy, hindlimb pseudofracture and muscle crush injury combined with hemorrhagic shock and resuscitation with either lactated ringers (LR) or FFP. **(B)** Volume of shed blood between LR and FFP groups compared with total volume of resuscitation. **(C)** Mean arterial pressure (MAP) measured at baseline and post-resuscitation. (D) Total white blood cell (WBC) count, **(E)** platelet count, **(F)** RBC count, and **(G)** hematocrit at 24 hours after resuscitation. (H) Heatmap of circulating inflammatory cytokines. **(I, J)** Representative images of pulmonary tissues from Sham, PT/HS-LR and PT/HS-FFP mice at 24 hours post-injury. The number of infiltrating **(I)** polymorphonuclear (PMN) leukocytes and **(J)** macrophages was quantified in relative to the total number of cells in each section. All scale bars = 50 μm. Data reported as mean ± standard error. \**p* < 0.05, \*\**p* < 0.01, \*\*\**p* < 0.001, \*\*\*\**p* < 0.0001, ns = not significant based on ANOVA and Fisher LSD post hoc test.

LR-resuscitated mice demonstrated increased pulmonary vascular permeability relative to sham mice, a pathologic finding that was not observed in the FFP group (**Figure 2A, B**). Interestingly, circulating levels of Angpt-2 were significantly elevated in both LR- and FFP-resuscitated PT/HS mice compared to shams at 24 hours (**Figure 2C**), indicative of pathological EC activation in both treatment groups. No significant differences were observed between LR and FFP groups in circulating levels of other markers of EC activation, including soluble ICAM-1, VEGF-A or VEGFR2 (**Figure 2D-F**).

**Figure 2.**
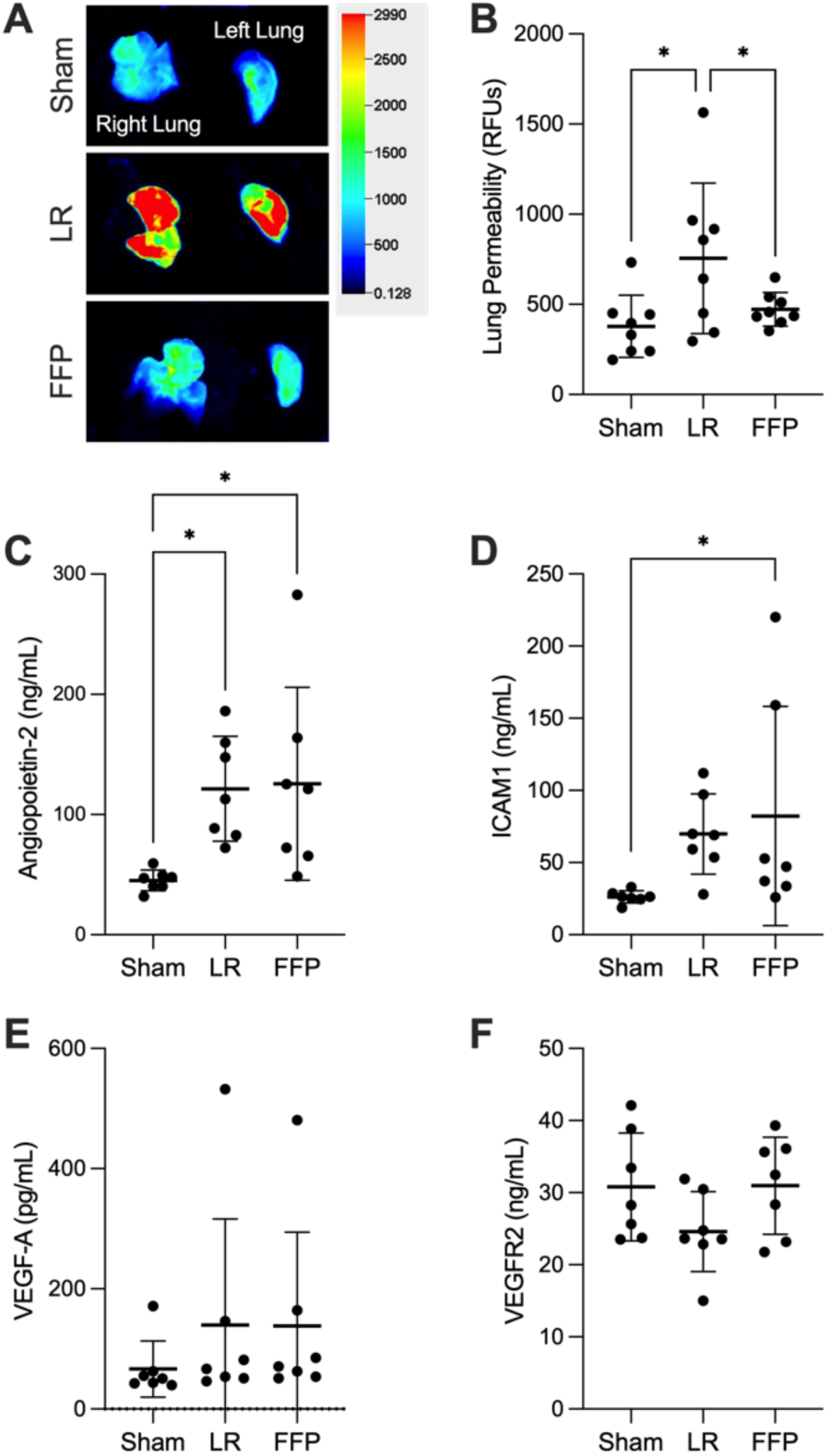
Fresh frozen plasma (FFP) decreases pulmonary vascular permeability after polytrauma and hemorrhagic shock (PT/HS). Pulmonary vascular leak was assessed based on extravasation of fluorescently-labeled dextran (70 kDa), followed by imaging and quantification. **(A)** Representative fluorescence images of lung tissue from sham, PT/HS-LR and PT/HS-FFP groups reflect the extent of dextran extravasation as a readout of vascular permeability. Heatmap colors indicate the magnitude of extravasated dextran, with red denoting greater leakage and blue indicating reduced permeability. **(B)** Pulmonary permeability was quantified as average fluorescent signal intensity (RFUs). **(C-F)** Plasma concentrations of angiopoietin-2, (D) ICAM, (E) VEGF-A, and (F) VEGFR2. \**p* < 0.05, based on ANOVA and Fisher LSD post hoc test.

### FFP maintains pulmonary eGC expression

Plasma levels of eGC biomarkers differed significantly between LR- and FFP-treated mice (**Figure 3A-D**), highlighting the effects of plasma-based resuscitation on eGC integrity. Specifically, circulating Sdc1 and HA levels increased after LR resuscitation, reflective of increased eGC damage, whereas FFP treatment was associated with increased levels of Sdc4, a HSPG involved in regulating endothelial pro-inflammatory signaling and tissue repair.^30–34^ In contrast, circulating glypican-1 levels were reduced in FFP mice compared to sham controls. WGA staining and TEM were used to visually assess pulmonary eGC expression after PT/HS and resuscitation. WGA signal intensity was diminished in LR-resuscitated mice compared to both sham and FFP-treated groups, indicating disruption of glycocalyx integrity (**Figure 4E**). TEM provided ultrastructural evidence of endothelial-specific glycocalyx expression, revealing a marked reduction of eGC in the pulmonary microvasculature of LR-resuscitated mice compared to sham controls (**Figure 4F**). In contrast, resuscitation with FFP enhanced eGC expression, as demonstrated by the presence of dense, negatively charged eGC aggregates. Together, these findings support the ability of FFP to preserve eGC expression and maintain vascular integrity following PT/HS.

**Figure 3.**
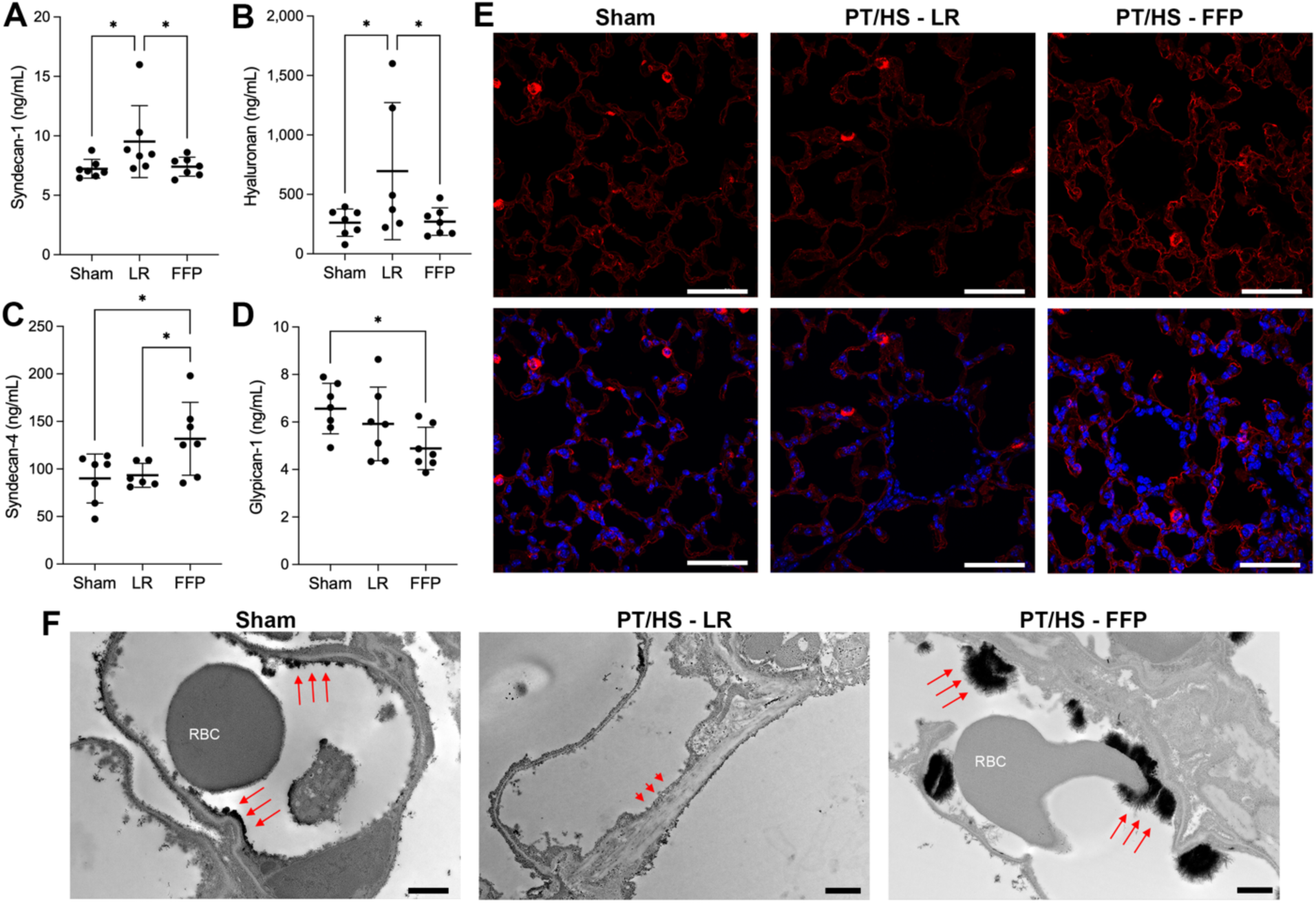
Fresh frozen plasma (FFP) enhances pulmonary endothelial glycocalyx expression after polytrauma and hemorrhagic shock (PT/HS). Plasma concentrations of **(A)** syndecan-1, **(B)** hyaluronan, **(C)** syndecan-4, and **(D)** glypican-1 and measured at 24 hours after sham procedure or PT/HS and resuscitation. \**p* < 0.05, based on ANOVA and Fisher LSD post hoc test. **(E)** Representative images of glycocalyx expression at 24 hours after injury, based on immunofluorescence staining using wheat germ agglutinin (red) and DAPI (blue). Scale bars = 50 μm. **(F)** Representative transmission electron micrographs of the glycocalyx lining the luminal surface of pulmonary endothelial cells as indicated by an electron-dense layer (long red arrows) that is notably absent in PT/HS-LR mice (short arrowheads). Scale bars = 1 μm.

**Figure 4.**
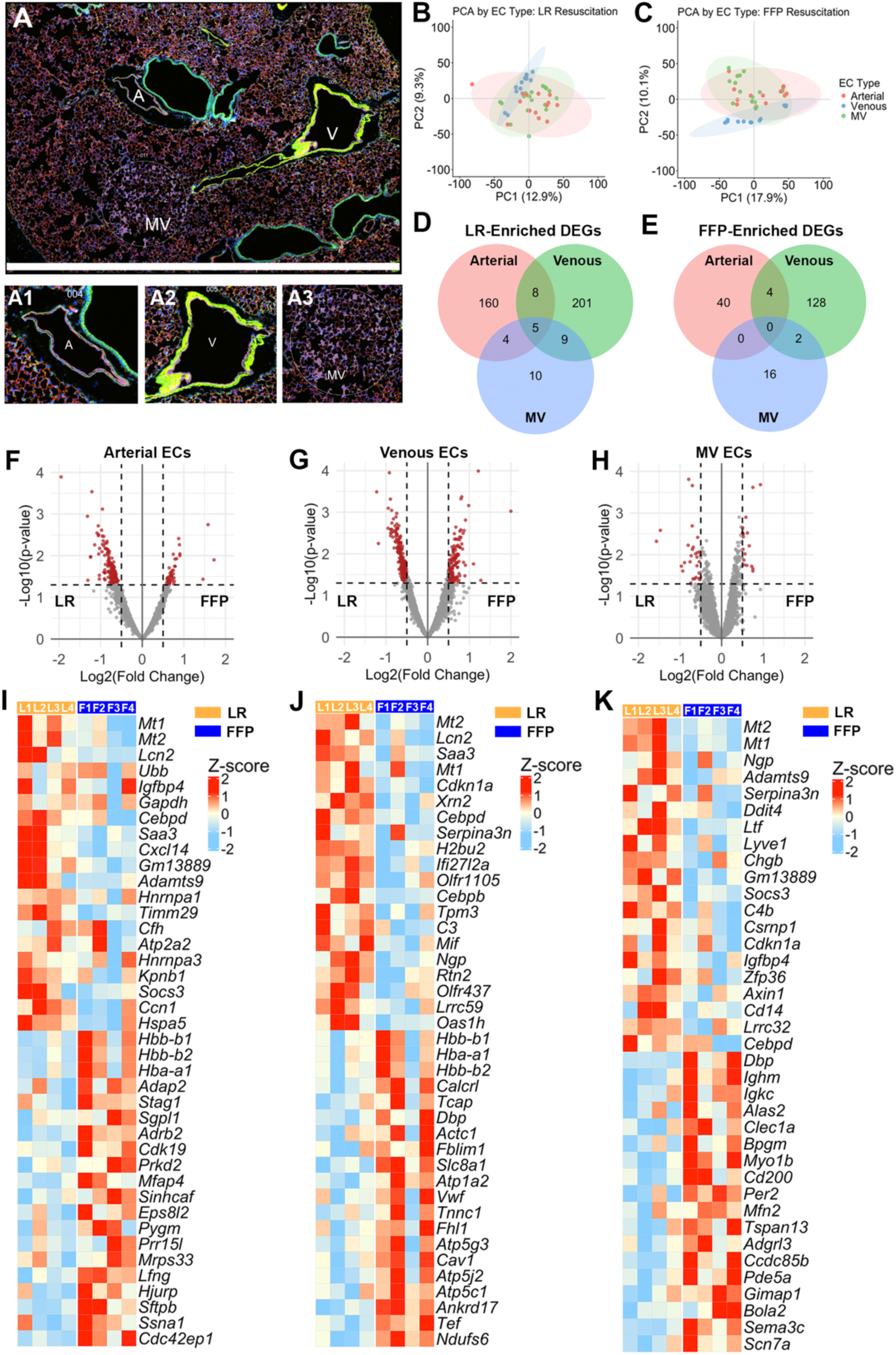
Resuscitation with fresh frozen plasma (FFP) elicits distinct, cell-type specific transcriptional responses across pulmonary arterial, venous and microvascular (MV) endothelial cells. **(A)** Representative epifluorescence image of mouse lung cross section captured on the GeoMx platform with insets demonstrating examples of regions of interest (ROI) from pulmonary arterial (A1), venous (A2), and microvascular endothelia (A3). **(B,C)** Principle component analyses of the ROIs from lactated Ringer’s (LR)- and FFP-treated mice. **(D,E)** Venn diagrams indicating the number of unique and overlapping differentially expressed genes enriched in LR- and FFP-treated mice, specific to each EC type. **(F-H)** Volcano plots depicting differential gene expression between arterial, venous and MV ECs after resuscitation with FFP (positive log2 fold change) or LR (negative log2 fold change). Red genes meet thresholds of p ≤ ; 1 × 10^−3^ and log2 fold change ≥ |1|. **(I-K)** Heatmaps depicting the top 40 differentially expressed genes enriched in **(I)** arterial, **(J)** venous, and **(K)** MV ECs after LR or FFP resuscitation. Each treatment group contains at least n = 3 EC-specific RNA samples from n = 4 mice per treatment condition. Colors represent average gene expression z-score for each biological replicate (L1-L4 and F1-F4), with red corresponding to upregulated and blue corresponding with downregulated expression.

### Plasma resuscitation effects distinct transcriptional programs in pulmonary arterial, venous and microvascular ECs after PT/HS

To gain insight into the vasculoprotective mechanisms associated with FFP resuscitation, spatial transcriptomic analysis of was performed in pulmonary arterial, venous, and MV ECs at 24 hours after PT/HS and resuscitation with FFP compared to LR (**Figure 4A-C**). Between the FFP and LR groups, a total of 221 differentially expressed genes (DEGs) were observed in arterial ECs, 357 DEGs in venous ECs and 46 DEGs in MV ECs (**Figure 4D-H**). Within each treatment group, there was minimal overlap in differential gene expression between arterial, venous and MV ECs (**Figure 4D-E**), indicating unique vessel-specific responses to resuscitation.

To visualize the most prominent transcriptional changes in arterial, venous, and MV ECs, heatmaps were generated to display the top 20 significant genes enriched in the FFP and LR groups, ranked by log-fold change (**Figure 4I-K**). Among the top genes that were differentially regulated, the metallothionein genes *Mt1* and *Mt2* were enriched in all EC types following LR resuscitation relative to FFP, likely indicative of exacerbated oxidative stress caused by crystalloid solutions and the activation of cytoprotective and immune regulatory pathways to promote cell survival after injury.^35,36^ FFP resuscitation resulted in enrichment of genes in arterial ECs that are associated with cytoskeletal remodeling (*Eps8l2*, *Cdc42ep1*),^37,38^ structural repair (*Hjurp*, *Ssna1*)^39,40^ and metabolic adaptation (*Mrps33*, *Pygm*).^41,42^ In venous ECs, *Ndufs6, Atp5g3*, *Atp5j2*, *Atp5c1*, and *Atp1a2* were among the top DEGs enriched after FFP resuscitation, supportive of elevated mitrochondrial respiration and increased ATP synthase production.^43,44^ Finally, in MV ECs, FFP resuscitation resulted in a mixture of gene signals related to mitochondrial function (*Mfn2*),^45^ circadian regulation of gene expression and metabolic cycles (*Dbp*, *Per2*),^46^ and genes such as *Myo1b*, *Ccdc85b*, *Tspan13*, *Adgrl3*, *Sema3c*, *Scn7a*, *Pde5a* that converge on adhesion and actin-myosin cytoskeletal remodeling programs.^47–49^

Based on our findings that FFP resuscitation was associated with preserved eGC integrity (**Figure 3**), we evaluated the expression of various genes involved in the biosynthesis, remodeling and maintenance of the eGC, including those that regulate HSPG synthesis, GAG chain polymerization, sulfation patterning of heparan sulfate, and protease- and enzyme-mediated eGC degradation/remodeling. Heatmap visualization of eGC-related genes revealed no dominant global clustering of genes across treatment groups, reflecting heterogenous expression profiles and variability across individual genes, EC types and mice (**Supplemental Figure 2**).

### FFP promotes EC signaling associated with tissue repair and enhanced cellular bioenergetics

Next, we performed GSEA-based pathway analysis to further explore the mechanistic impact of FFP on the pulmonary vasculature. Venous ECs were selected for these analyses due to the limited number of DEGs in arterial and MV ECs. Based on predefined thresholds, 238 and 33 Reactome pathways were enriched in ECs following FFP resuscitation and 34 GO:Biologic Process and 6 Reactome pathways were enriched after LR resuscitation (**Supplemental Table 1**). To derive mechanistic insight into EC responses to trauma based on the GO and GSEA results, subsequent analyses focused on non-redundant pathways with high gene overlap and direct relevance to 1) stress and ennvironmental response, 2) regulation of cellular processes, 3) signaling and kinase regulation, 4) vascular development and regulation, 5) metabolism and biosynthesis, and 6) stem cell biology (**Supplemental Table 1**). The top 3 GO:Biologic Process pathways within each category, based on the number of overlapping genes, and all Reactome pathways are shown in **Figure 5**.

**Figure 5.**
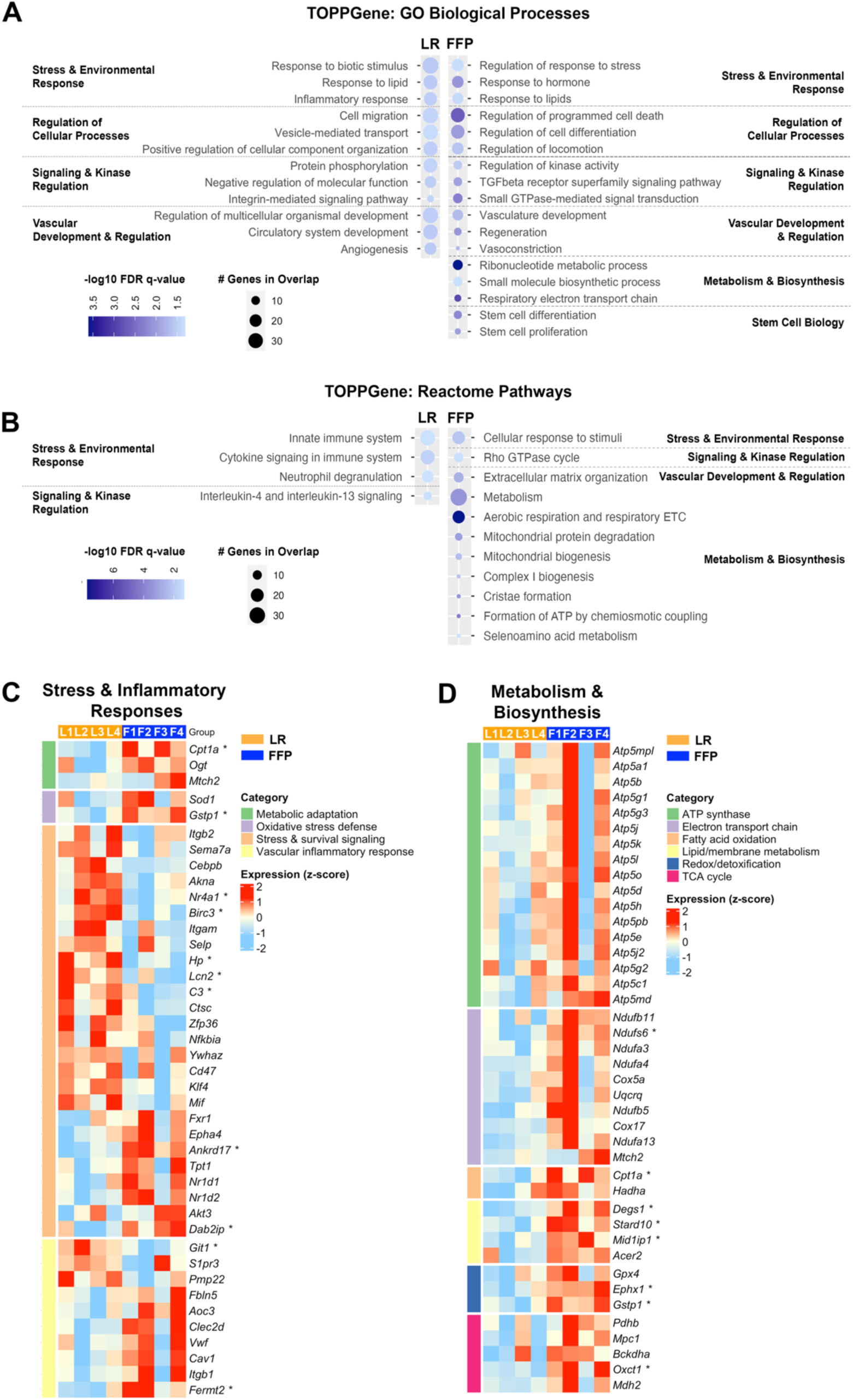
Gene set enrichment analysis (GSEA) in pulmonary venous endothelial cells (EC). GSEA was performed using TOPPGene with **(A)** Gene Ontology (GO): Biological Process and **(B)** Reactome pathway datasets. Pathways with ≥ 5 overlapping gene counts and false discovery rate (FDR)-adjusted *p* value (*q* < 0.05) were categorized according to their relevance to vascular and/or trauma biology. The figure displays the top 3 GO: Biological Process pathways and all identified Reactome pathways within each category. **(C,D)** Specific genes associated with GO and Reactome pathways governing cellular stress, inflammatory and metabolic responses to injury are presented in heatmap plots, categorized according to relevance to general cellular and biologic processes. Heatmap colors represent average gene expression z-score for each biological replicate (L1-L4 and F1-F4), with red corresponding to upregulated and blue corresponding with downregulated expression. For each gene, differences between groups were evaluated using Welch’s t-test on unscaled tissue-level expression values (\**p* < 0.05).

Whereas LR resuscitation was predominantly associated with proinflammatory immune responses, FFP resuscitation led to an enrichment in biological pathways associated with adaptive cell signaling responses, extracellular matrix remodeling, protection against programmed cell death and mitochondrial energy metabolism (**Figure 5A, B**). We also observed GO terms associated with tissue regeneration, including stem cell development and proliferation in ECs from FFP-resuscitated mice, indicating that FFP may promote regenerative mechanisms to repair endothelial damage after injury. In line with these findings, the GO:Biologic pathway encoding the TGF-β receptor signaling pathway, an established promoter of wound healing and tissue repair ^50,51^, was significantly enriched in FFP-treated ECs (**Figure 5A**). Moreover, we observed robust enhancement of mitochondrial and metabolic Reactome pathways after FFP resuscitation (**Figure 5B**), indicating the potential for FFP to promote mitochondrial biogenesis after injury, which could help to mitigate oxidative stress caused by injury in addition to supporting increased energy demands required for endothelial repair.

To gain additional insight, genes associated with stress, inflammatory and metabolic response pathways were plotted in heatmaps (**Figure 5C, D**). Categorical grouping of genes highlighted significant associations between FFP and a transcriptional signature consistent with mitochondrial stress adaptation (*Cpta1*),^52^ oxidative stress defense (*Gstp1*),^53^ and cell cycle progression (*Ankrd17*),^54^ alongside a shift towards cell-ECM stabilization (*Fermt2*)^55^ and reduced migratory signaling (*Git1*)^56^ (**Figure 5C**). Conversely, after LR resuscitation, gene signatures reflected robust innate and acute-phase activation (*Birc3*, *Lcn2*, *C3*)^57–59^ with reduced counter-regulation of NF-κB-mediated inflammatory signaling (*Dab2ip*).^60^ Furthermore, transcriptional changes caused by FFP suggested overall enhanced mitochondrial bioenergetics compared to LR resuscitation, including enrichment of genes associated with fatty acid oxidation, TCA flux, electron transport chain activity and ATP synthase (**Figure 5D**).

### FFP resuscitation increases mitochondrial content in EC

To validate transcriptional responses of pulmonary venous EC to PT/HS and FFP resuscitation, we performed histological assessments of mitochondrial content in the pulmonary vasculature of FFP- and LR-resuscitated mice using TOM20, an outer mitochondrial membrane import receptor widely used as a marker of mitochondrial mass that correlates with oxidative capacity.^61^ ECs from FFP-resuscitated mice had signficantly greater expression of TOM20 than LR-treated mice (**Figure 6A,B**), supporting results from our GSEA analyses that FFP resuscitation is associated with mitochondrial biogenesis and related functional pathways. Similarly, *in vitro* treatment of EC with 30% FFP for 6 hours also resulted in increased TOM20 expression compared to LR-treated cells, confirming that FFP enhances endothelial-specific mitochondrial content in primary human lung EC (**Figure 6C,D**).

**Figure 6.**
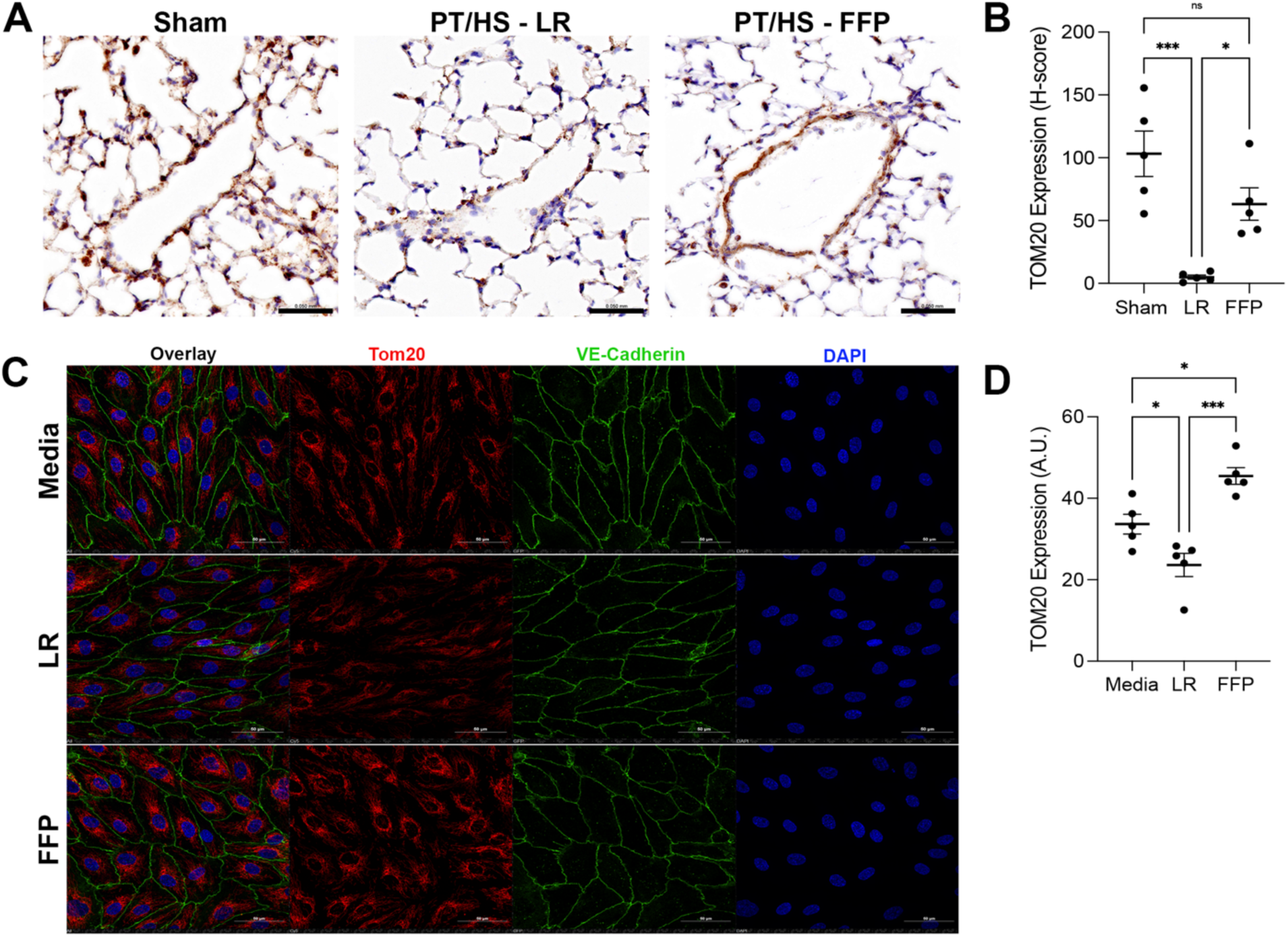
Mitochondrial content is enhanced in pulmonary endothelial cells (EC) after treatment with fresh frozen plasma (FFP) compared to lactated Ringer’s (LR). **(A)** Immunohistology images of translocase of outer membrane mitochondria 20 (TOM20) expression in pulmonary EC at 24 hours after sham procedure or polytrauma-hemorrhagic shock (PT/HS) and resuscitation with either FFP or LR. **(B)** The histoscore (H-score) was used to semi-quantify TOM20 staining intensity in n = 5 randomly selected venous structures for each tissue. **(C)** Immunofluorescence images of TOM20 in human lung EC treated with 30% (v/v) FFP or LR for 6 hours. Vascular endothelial (VE)-cadherin and DAPI were used to visualize the cellular monolayer and nuclei, respectively. **(D)** TOM20 staining intensity was quantified in ImageJ and expressed as mean gray value in arbitrary units (A.U.). All scale bars = 50 μm. Data reported as mean ± standard error. \**p* < 0.05, \*\**p* < 0.01, \*\*\**p* < 0.001, ns = not significant based on ANOVA and Fisher LSD post hoc test.

## Discussion

In this study we investigated the mechanistic effects of FFP-based resuscitation after polytrauma on vascular EC integrity compared to resuscitation with LR crystalloid and sham controls. We demonstrated that 24 hours after PT/HS FFP administration significantly decreased systemic levels of inflammatory biomarkers including IL-6, IL-1α, and TNF-α compared to LR-treated mice. We also noted a significant reduction in lung permeability in FFP-treated mice compared to LR controls and found that circulating biomarker levels of syndecan-1, glypican-1, and hyaluronan were significantly reduced in the FFP-treated cohort compared to controls. Visually, FFP resuscitation also resulted in more robust eGC expression within the pulmonary vasculature, indicating a possible regeneration and stabilization of the EC membrane. Utilizing spatial transcriptomics, we examined gene expression within pulmonary MV, arterial and venous EC further confirming pulmonary EC stability after FFP administration based on an enrichment of genes involved in mitochondrial activation and aerobic respiration, compared to genes related to pro-inflammatory and innate immune responses that were associated with LR resuscitation.

Modern resuscitation strategies emphasize early administration of plasma in hemorrhaging patients, which mitigates coagulopathy, reduces transfusion requirements, and improves mortality. However, the mechanistic underpinnings for these clinical benefits have not been fully elucidated. Previous *in vitro* work has shown that FFP can directly modulate endothelial expression of adhesion molecules to mitigate neutrophil-EC interactions,^62^ and preclinically, resuscitation with FFP restores vascular integrity, at least in part by, promoting eGC expression and decreasing pulmonary endothelial hyperpermeability.^13,29,63,64^ Our current results replicate these prior findings, confirming the beneficial effect of FFP on endothelial integrity at 24 hours after injury. Most preclinical models of hemorrhagic shock and resuscitation examine the acute effects of FFP on the eGC, excluding analyses at later timepoints after injury. Our study utilized a more severe model of polytraumatic and hemorrhagic injuries resulting in sustained endotheliopathy for 24 hours after LR resuscitation, which was not observed in other murine models.^26^ In our study, inclusion of peripheral injury via hindlimb muscle crush and pseudo-fracture likely contributed to sustained systemic endothelial dysfunction. We also included large volume crystalloid resuscitation with LR, which although is not the current clinical standard for treating hemorrhagic shock, induces severe endotheliopathy and organ injury—effects that our data show can be counterbalanced by pressure-guided resuscitation with FFP.

This study provides new insight into the endothelial-specific transcriptional landscape of the pulmonary vasculature after resuscitation with FFP, highlighting many signaling pathways that may govern vasculoprotective responses to FFP. Compared to FFP-treated mice, we identified an enrichment of metallothionein genes following LR resuscitation indicative of increased oxidative stress. It is well known that hemorrhagic shock contributes to a significant shift in cell energy metabolism, from aerobic to anaerobic metabolism contributing to a profound metabolic acidosis. Once the acidotic insult has occurred, cell cycle signaling, and receptor activation may become impaired along with an increase in cell death.^65^ D’Alessandro et al.^66^ found that in a rodent model of hemorrhagic shock, FFP-based resuscitation corrected hyperlactatemia, potentially fueling mitochondrial metabolism through utilization of alternative metabolic pathways not noted in normal saline resuscitated rodents, for example via the TCA cycle as evidenced by notable elevations in glutamine catabolites. Based upon these results, one could infer that the vasculoprotective properties of FFP resuscitation may be related to a resolution of intracellular pH lactic acidosis that, in turn, contributes to structural stability of the eGC and/or regulation of eGC-protein interactions that mediate vascular repair and homeostasis. Although these potential associations between metabolic acidosis and eGC integrity are intriguing, further research is needed to more clearly define causal mechanisms and the resultant implications on vascular repair and homeostasis after trauma.

Another possible mechanistic benefit of FFP-based resuscitation is through the promotion of mitochondrial energy metabolism and biogenesis after traumatic injury, evidenced by preponderance of metabolic-related pathways identified in our untargeted genomic analysis. More specifically, we identified that genes *Atp5g3, Atp5j2, Atp5c1* were generally elevated after FFP resuscitation. These genes are known to encode ATP synthase of complex V within the mitochondria which is the final complex of oxidative phosphorylation.^67^ Futhermore, TOM20 immunostaining provides further evidence of FFP’s ability to preserve mitochondrial content compared to LR, findings that were also confirmed in primary human ECs. Based upon these findings, FFP-based resucitation may promote mitochondrial biogenesis and enhance overall mitochondrial functions related to cellular ATP production, contributing to improved EC homeostasis and reduced end organ dysfunction after traumatic insult. Villarroel et al.^68^ demonstrated that exposure of peripheral blood mononuclear cells isolated from traumatically injured rats to healthy plasma promoted mitochondrial recovery through an upregulation of oxidative phosphorylation and ATP production. Likewise, Hwabejire et al.^69^ found a significant improvement in mitochondrial cell membrane stabilization after FFP resuscitation in a porcine model of PT/HS and TBI, along with a notable increase in mitochondrial cell function as measured through pyruvate dehydrogenase activity compared to normal saline controls. Researchers concluded that FFP-based resuscitation may reduce overall mitochondrial apoptosis through mitochondrial cell wall stabilization leading to an improvement in cellular bioenergetics.^69^ Alongside our current EC-specific findings, these studies provide compelling evidence that mitochondrial function is central to FFP-mediated endothelial stabilization that governs vascular integrity and end organ (dys)function after injury.

With regards to eGC expression, our data did not reveal robust differences between FFP and LR mice in gene expression related to proteoglycan and GAG biosynthesis, modification or degradation/remodeling. The absence of a strong, unified transcriptional signature of eGC synthesis or stabilization may reflect the timing of the transcriptomic analysis, which may not coincide with peak or coordinated transcriptional activation of eGC repair pathways following FFP and LR resuscitation. These findings may also suggest that preservation by FFP is mediated through post-transcriptional, enzymatic, or biophysical mechanisms rather than sustained large-scale changes in gene expression.

This study has important limitations. First, our analyses were performed at 24 hours after resuscitation, limiting insight into the acute effects of FFP on the vascular endothelium and eGC expression. Instead, our findings describe the more sustained impact of FFP on pulmonary ECs in relation to endotheliopathy that is linked to systemic and localized tissue inflammation. Second, endotheliopathy in other organ systems known to be impacted after PT/HS, such as the kidneys, liver and mesentery, was not evaluated in this study, and additional work is needed to determine whether the transcriptional responses to FFP identified within the pulmonary vasculature extend to these other organs. Third, we only utilized male mice within a defined age window in this study. Future investigations with both male and female mice of varying age are warranted to understand if age and/or systemic hormonal differences impact endothelial stabilization after FFP resuscitation.

In conclusion, goal-directed plasma resuscitation after PT/HS decreases systemic inflammation, pulmonary endovascular leak and recapitulates the pulmonary eGC. This study provides novel insight into pulmonary endothelial-specific transcriptional responses to FFP resuscitation that may link post-injury eGC integrity with mitochondrial bioenergetics and metabolism, revealing potential vasculoprotective mechanisms by which FFP alleviates multiorgan failure.

## Acknowledgements

This project was supported by National Institutes of Health R35GM137958 to JRR. TEM imaging was supported by the UAB High Resolution Imaging Facility.

---

**Supplemental Figure 1.**
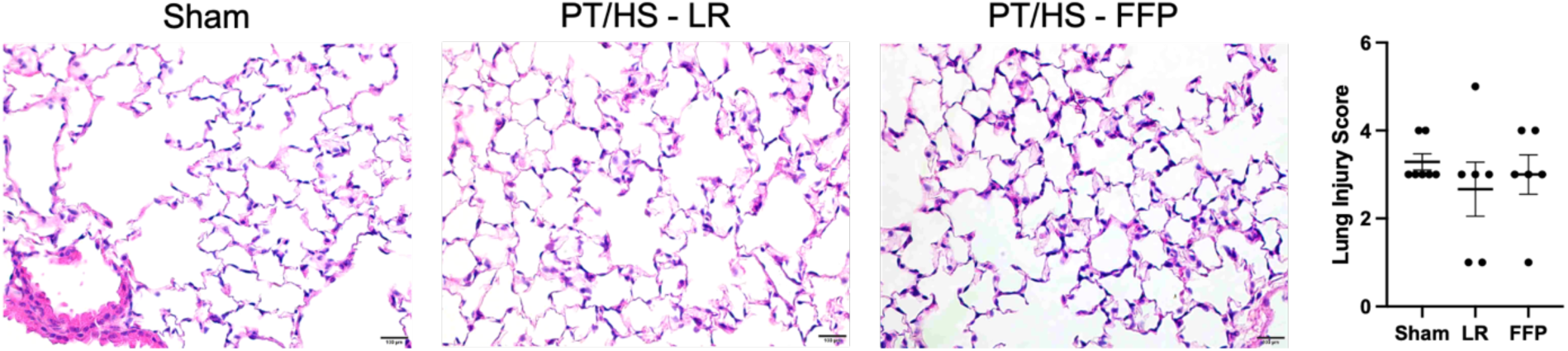
Histologic lung injury 24 hours after sham procedure or polytrauma-hemorrhagic shock (PT/HS) and resuscitation with either lactated Ringer’s (LR) or fresh frozen plasma (FFP). H&E stained lung slices were evaluated by a blinded pathologist and scored for histologic injury. Representative images at 20x are provided. N=6-7 per group. Scale bars = 100 μm. Data reported as mean ± standard error.

**Supplemental Figure 2.**
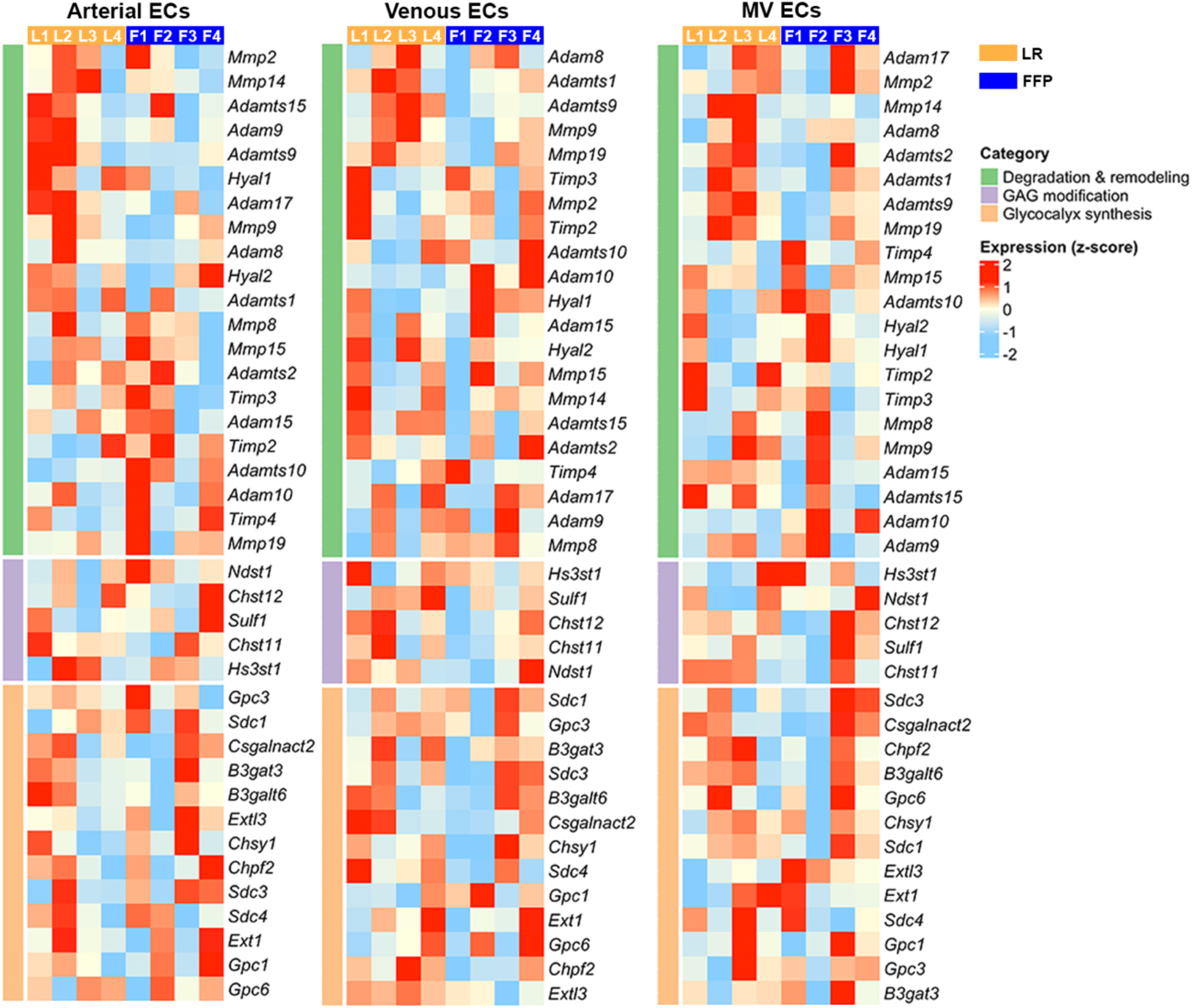
Heatmaps showing expression of genes related to endothelial glycocalyx synthesis, degradation/remodeling and sulfation across pulmonary arterial, venous and microvascular endothelial cells.

